# Target enrichment improves phylogenetic resolution in the genus *Zanthoxylum* (Rutaceae) and indicates both incomplete lineage sorting and hybridization events

**DOI:** 10.1101/2021.04.12.439519

**Authors:** Niklas Reichelt, Jun Wen, Claudia Pätzold, Marc S Appelhans

## Abstract

**Background and aims:** *Zanthoxylum* L. is the only pantropical genus within Rutaceae, with a few species native to temperate eastern Asia and North America. Efforts using Sanger sequencing failed to resolve the backbone phylogeny of *Zanthoxylum*. In this study, we employed target enrichment high-throughput sequencing to improve resolution. Gene trees were examined for concordance and sectional classifications of *Zanthoxylum* were evaluated. Off-target reads were investigated to identify putative single-copy markers for bait refinement, and low-copy markers for evidence of putative hybridization events.

**Methods:** We developed a custom bait set for target enrichment of 745 exons in *Zanthoxylum* and applied it to 45 *Zanthoxylum* species and one *Tetradium* species as the outgroup. Illumina reads were processed via the HybPhyloMaker pipeline. Phylogenetic inferences were conducted using coalescent and concatenated methods. Concordance was assessed using quartet sampling. Off-target reads were assembled and putative single- and low-copy genes were extracted. Additional phylogenetic analyses were performed based on these alignments.

**Key results:** Four major clades are supported within *Zanthoxylum*: the African clade, the *Z. asiaticum* clade, the Asian-Pacific-Australian clade, and the American-eastern Asian clade. While overall support has improved, regions of conflict are similar to those previously observed. Gene tree discordances indicate a hybridization event in the ancestor of the Hawaiian lineage, and incomplete lineage sorting for the American backbone. Off-target putative single-copy genes largely confirm on-target results, and putative low-copy genes provide additional evidence for hybridization in the Hawaiian lineage. Only two of the five sections of *Zanthoxylum* are resolved as monophyletic.

**Conclusion:** Target enrichment is suitable to assess phylogenetic relationships in *Zanthoxylum*. Our phylogenetic analyses reveal that current sectional classifications need revision. Quartet tree concordance indicates several instances of reticulate evolution. Off-target reads are proven useful to identify additional phylogenetically informative regions for bait refinement or gene tree based approaches.

## Introduction

With the advance of Next-Generation-Sequencing (NGS) approaches in systematics, hitherto recalcitrant phylogenetic relationships, *i.e*. rapid radiations (Welch et al., 2016) or deep divergences (Zeng et al., 2014), can be tackled with increasingly large datasets at steadily decreasing costs (Straub et al., 2012). The majority of the NGS approaches in systematics aim to achieve a reduced representation of the genome to exclude regions with low phylogenetic signal and reduce computational complexity (Hörandl and Appelhans, 2015; Zimmer and Wen, 2015). Different methods have emerged, varying in applicability at different taxonomic levels and with regards to sample conservation. For target enrichment methods, regions of interest are captured and isolated via biotinylated RNA baits designed using reference data (Lemmon and Lemmon 2012; Weitemier et al., 2014). One major advantage of target enrichment methods is the applicability to both herbarium, silica-gel preserved and fresh material (Villaverde et al., 2018), while obtaining and analyzing high quality genomic or transcriptomic sequence data from target or closely related species to identify orthologous and phylogenetically informative regions in advance is often the greatest challenge (Twyford and Ennos, 2012). However, this disadvantage is mediated by the increasing availability of transcriptomic and genomic data across angiosperm plant families. The mapping rates of reads to baits are often in the range of 60-80%, but rates of 20% or less have also been reported (Schmickl et al., 2016; Soto Gomez et al., 2019; Tomasello et al., 2020). Thus, off-target reads may serve as a useful resource in target enrichment approaches. While a fraction of the off-target reads has been frequently utilized to assemble plastid genes or genomes as a “by-product” (e.g., Weitemier et al., 2014; Ma et al., 2021), the remaining off-target reads may be utilized further. They might be used to assemble additional, un-targeted single or low-copy regions, which in turn might be used to expand the existing dataset, to refine the bait set for further approaches, to investigate reticulate evolution or evolution of gene families amongst others.

*Zanthoxylum* L. (prickly ash, yellowwood) belongs to subfamily Amyridoideae (Morton and Telmer, 2014) and represents the second largest genus within Rutaceae with about 225 currently accepted species (Kubitzki et al., 2011). It is distributed over all continents except Europe and Antarctica with biodiversity hotspots in the (sub-)tropics. A few species are adapted to a colder climate and are native to North America and temperate eastern and southern Asia (Reynel, 2017), where they have been used widely as spice (e.g., Sichuan pepper, sanshō pepper, timur) or herbal medicine (e.g., Lu et al. 2020). Most *Zanthoxylum* species can be easily recognized by thorny bosses on the trunk and branches, and spines may be found at a pseudo-stipular position (Weberling, 1970) and/or along the rachis of leaves or leaflets (Zhang et al., 2008). *Zanthoxylum* has an alternate phyllotaxis with punctate, estipulate and usually pinnate leaves. The plants are usually dioecious and the perianth may be homo- or heterochlamydeous. Seeds stay attached to the opening fruits (follicles) (Hartley, 1966; Kubitzki et al., 2011) and may be dispersed by birds (Silva et al., 2008; Guerrero and Tye, 2009), mammals (Muller-Landau et al., 2008), ants (Maschwitz et al., 1992; Reynel, 1995), or fish (Reys et al., 2009). Most species reproduce sexually (Kamiya et al., 2008; Costa et al., 2013), but apomixis via nucellar embryony has also been reported (Liu et al., 1987; Naumova, 1993). Due to a significant variation in the flower morphology of *Zanthoxylum*, Linnaeus (1753, 1759) differentiated between *Zanthoxylum* s.str. with homochlamydeous flowers, and *Fagara* L. with heterochlamydeous flowers. Brizicky (1962) hypothesized that the simple perianth in *Zanthoxylum* s.str. is a secondary condition that derived from the double perianth of *Fagara.* Today, both taxa are united in the morphologically diverse *Zanthoxylum* s.l. since *Zanthoxylum* s.str. is deeply nested within *Fagara* (Appelhans et al., 2018). The most recent taxonomic treatment based on morphological traits was published by Reynel (2017) and will be used herein. According to Reynel (2017), *Zanthoxylum* species formerly accounted to *Fagara* are ascribed into the pantropical section *Macqueria* and the American sections *Tobinia* and *Pterota*. Members of *Zanthoxylum* s.str. are divided into an American section *Zanthoxylum* and an Asian section *Sinensis*.

Phytochemical (Waterman, 2007) and DNA sequence data (Poon et al. 2007; Appelhans et al. 2018) have confirmed that *Zanthoxylum* is most closely related to *Tetradium* Lour., *Phellodendron* Rupr. from Asia, and *Fagaropsis* Mildbr. ex Siebenlist from Africa and Madagascar. The monotypic *Toddalia* Juss. was recently merged with *Zanthoxylum* (Appelhans et al., 2018) and shows a broad distributional range from tropical Africa and Madagascar to eastern and southeastern Asia. A rich fossil record is evident in Eocene Europe for all these genera except *Fagaropsis* (Chandler 1961; Gregor, 1989; Collinson et al., 2012). *Zanthoxylum* has been absent from Europe since the late Miocene to early Pliocene (Geissert et al., 1990) but spread over all other continents except Antarctica (*i.e*., Graham and Larzen, 1969; Jacobs and Kabuye, 1987; Tiffney, 1994; Kershaw and Bretherton, 2007). Recently, we conducted a first worldwide phylogenetic study of *Zanthoxylum* with 99 specimens comprising 54 species (Appelhans et al. 2018). However, based on only two nuclear and two plastid markers, several nodes in the backbone phylogeny remained unresolved, especially regarding the American and Pacific lineages. The Pacific *Zanthoxylum* lineage was resolved as monophyletic in the plastid dataset but polyphyletic in the nuclear dataset, possibly related to a previous hybridization event.

Here, target enrichment is applied in *Zanthoxylum.* We first design a bait set based on newly generated transcriptome data and test its suitability for phylogenetic reconstructions in the genus. The main goal of this study is to improve phylogenetic resolution regarding the main clades within the genus (Appelhans et al. 2018). The large quantity of sequence data will help evaluate whether the low resolution in previous Sanger sequencing studies (Appelhans et al., 2018) was due to a lack of informative characters or cases of reticulate evolution or incomplete lineage sorting. An additional goal is to test the most recent sectional classifications by Reynel (2017) using the phylogenetic framework. Finally, we aim to explore whether off-target reads can be used to identify additional informative regions that can be used in phylogenetic analyses and to improve and/or enlarge bait sets for future studies.

## Materials and methods

### a. RNA-Seq and bait design

We employed a “made-to-measure” bait design strategy (Kadlec et al., 2017) to increase bait specificity. Transcriptomic data of four *Zanthoxylum* accessions (representing three species) and three closely related outgroups served as foundation for bait design (Supplemental Tab. S1). Three transcriptomes were publicly available via the NCBI SRA archive, and four additional transcriptomes were generated in the course of this study (Supplemental Tab. S1). Young leaves of plants cultivated at Goettingen Botanical Garden were frozen in liquid nitrogen for RNA preservation. Total RNA was extracted using the RNeasy® Plant Mini Kit (Qiagen) as per the manufacturer’s instructions. Library preparation for Illumina sequencing was performed at the Transcriptome and Genome Analysis Laboratory Goettingen (TAL) using the TruSeq RNA Library Prep Kit v2 (Illumina, San Diego, CA, USA). Pooled libraries were run on an Illumina HiSeq 4000 to produce 50bp single end reads. Raw sequence data were trimmed using cutadapt v1.1.6 (Martin, 2011), removing adapter sequences with a minimum overlap of 10 bp and trimming read ends showing a PHRED score of less than 30. Trimmed reads with a remaining length of less than 35 bp were discarded. Trinity v2.5.1 (Grabherr et al., 2011; Haas et al., 2013) was used for *de novo* assembly of trimmed reads using default options with the exception of *max. memory*, which was set to 50GB. Identification of single copy orthologous loci was conducted as described in Tomasello et al. (2020) using a combination of MarkerMiner (Chamala er al., 2015) and custom python scripts (https://github.com/ClaudiaPaetzold/MarkerMinerFilter). Exons shorter than 120 bp were discarded. The variability between only *Zanthoxylum* sequence data was assessed and regions showing less than 0.5% or more than 15% variability were discarded. The obtained 745 exon sequences spanning 354 genes were further processed by Arbor Biosciences (myBaits®, Ann Arbor, MI, USA), which included masking, to produce a set of 20,000 80-mer baits.

### b. Sampling and DNA extraction

We sampled a total of 47 *Zanthoxylum* specimens representing 44 different species and one specimen of *Tetradium* (Tab. 1) as outgroup. All major distributional areas and sections according to Reynel (2017) are covered. Total DNA was extracted from herbarium or silica-dried material using a variation of the CTAB protocol by Doyle and Doyle (1987) or the DNeasy Plant Mini Kit (Qiagen, Hilden, Germany) following manufacturer’s instructions.

**Tab. 1.**
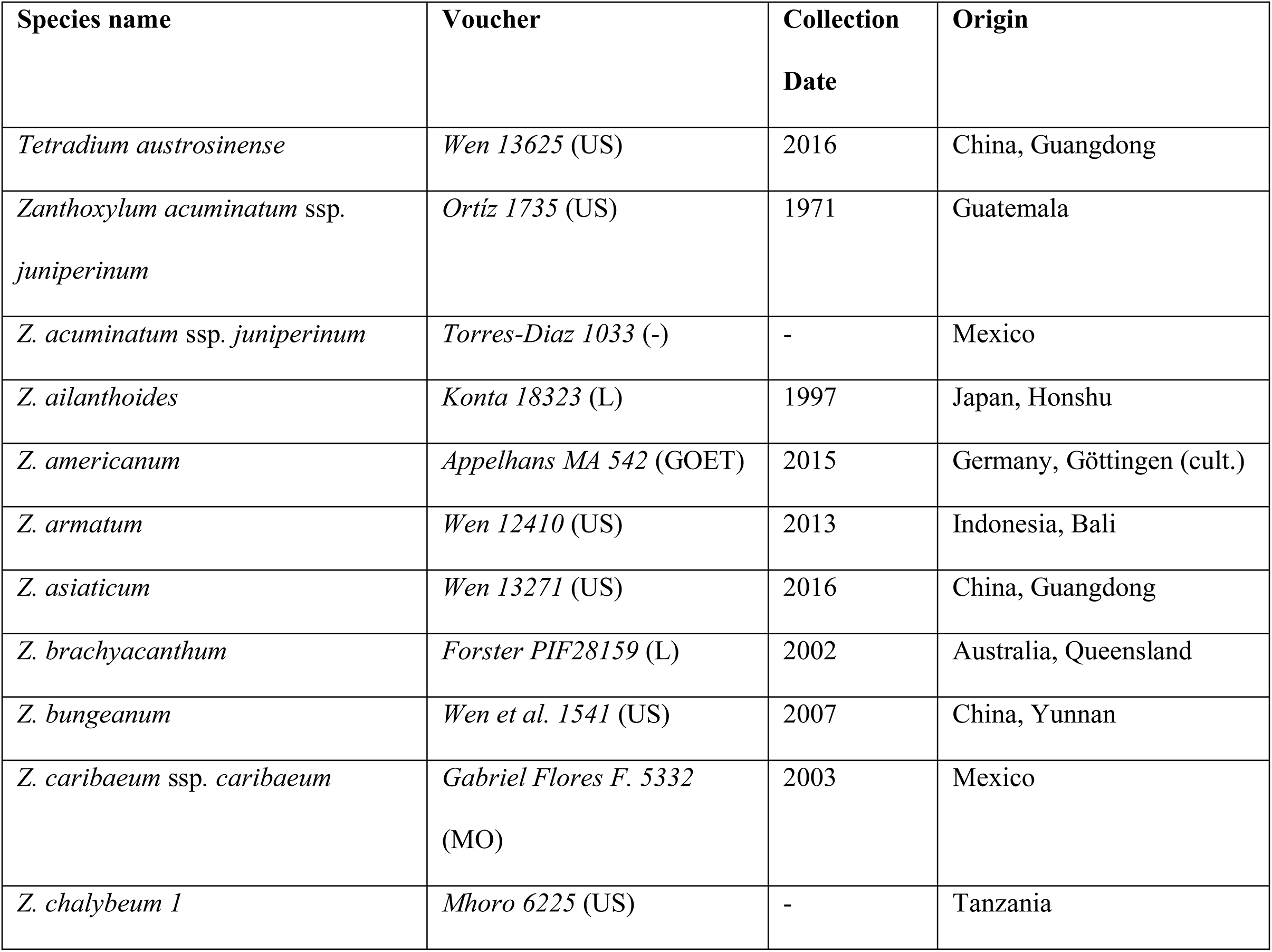
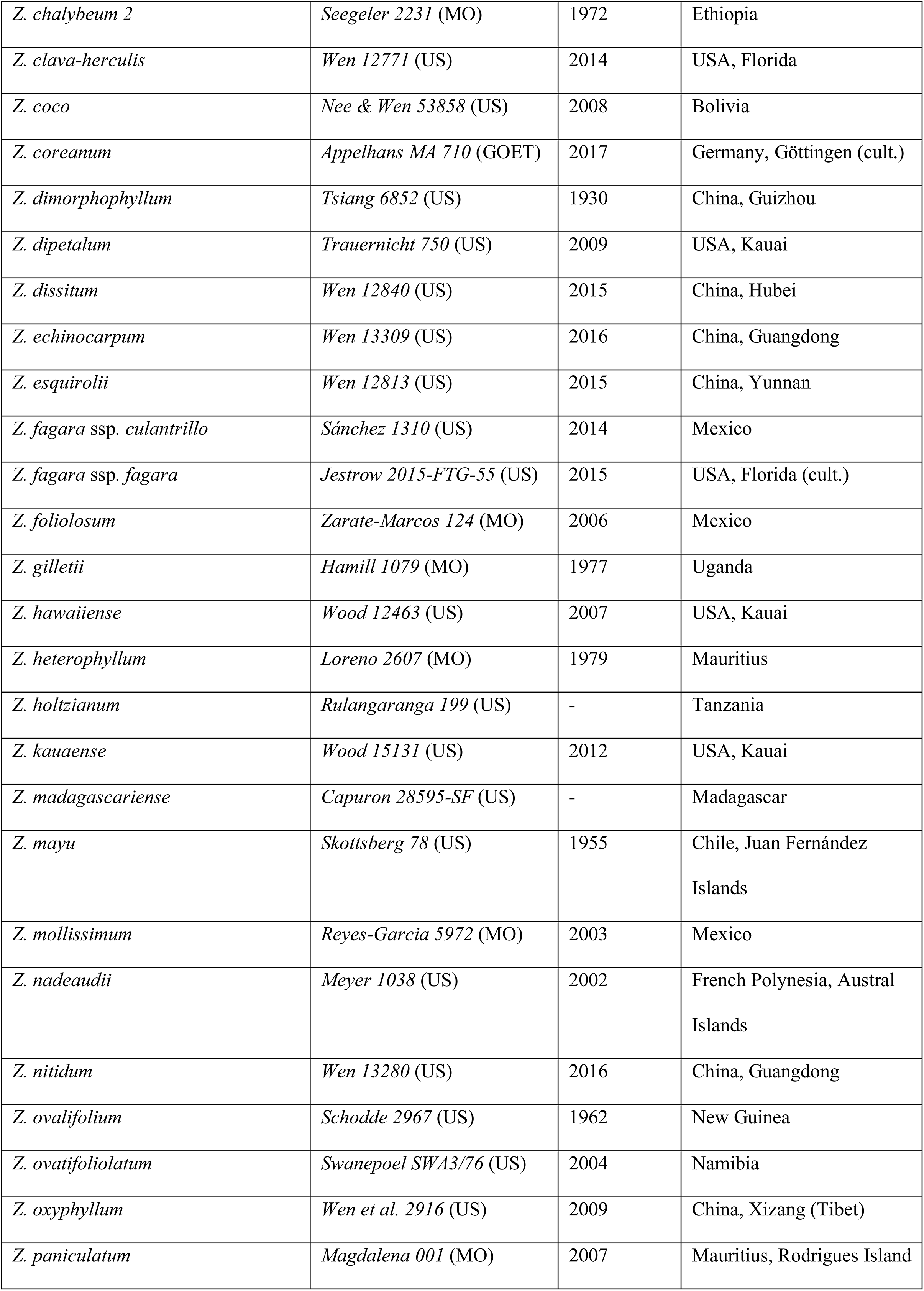
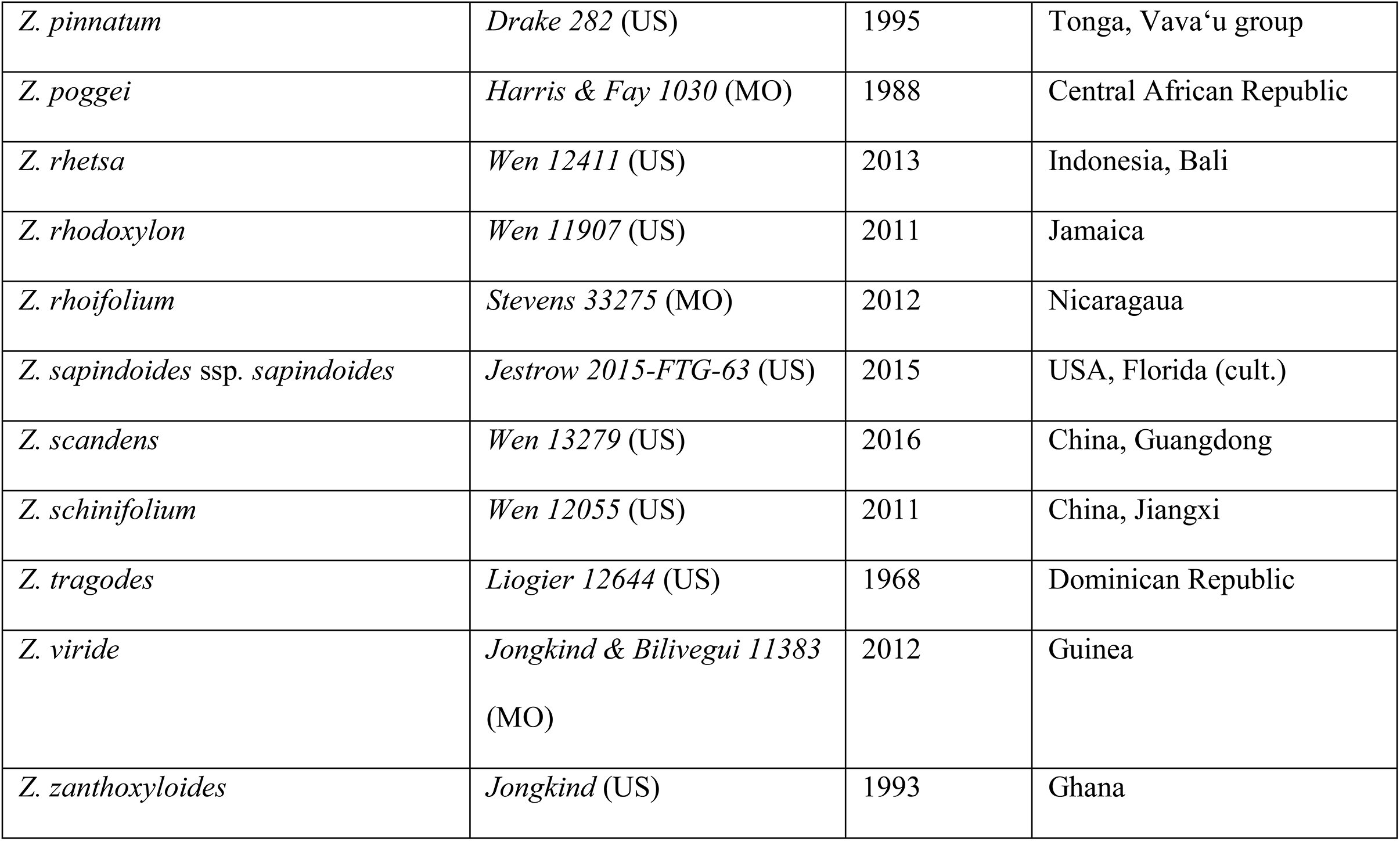
Specimens sampled for target enrichment and phylogenetic analysis including voucher information, date of collection and geographic region. *T.* = *Tetradium*. *Z.* = *Zanthoxylum*.

### c. Library Preparation for Target Enrichment and Sequencing

For each sample we used a Q800R sonicator (Qsonica; Newtown, CT, USA) to shear 800 ng of DNA to an approximate fragment size of 350 bp. Library preparation was conducted using the NEBNext® Ultra™ II DNA Library Prep Kit for Illumina® (New England Biolabs, Ipswich, MA, USA) with NEBNext Multiplex Oligos for Illumina® (Dual Index Primers Set 1) and AMPure XP magnetic beads (Beckman Coulter, Brea, CA, USA). DNA content and quality of indexed libraries were examined with a Qubit 4.0 using the high-sensitivity kit (ThermoFisher Scientific, Waltham, MA, USA) and a 1% agarose gel. Samples were pooled in pairs of two or four based on their respective quality, resulting in pools containing a total of 500 ng equimolar DNA. Hybridization of baits to pooled libraries was conducted according to the myBaits® - Hybridization Capture for Targeted NGS – Manual v4.01 (April 2018; Arbor Biosciences) with the exception of hybridization time, which was extended to 40 h. Captured DNA libraries were amplified using the KAPA HiFi HotStart Ready Mix (Hoffmann-La Roche, Basel, Switzerland) with the suggested maximum of 14 cycles. Enriched libraries were purified with AMPure XP magnetic beads and checked for quality with qPCR, using i5 and i7 Illuima TruSeq primers, and Bioanalyzer (Agilent Technologies, Santa Clara, CA, USA). Samples were sequenced on an Illumina HiSeq 4000 producing 2×150 bp paired-end reads (Illumina, San Diego, CA, USA) at Novogene (Sacramento, CA, USA).

### d. Data Analysis of Targeted Reads

The HybPhyloMaker pipeline (Fér and Schmickl, 2018) provided the bash scripts for all the steps from initial quality trimming of reads to species tree reconstruction. Trimmomatic v0.33 (Bolger et al., 2014) was used for adapter and quality trimming with thresholds set to a minimum of 65 bp read length and a minimum PHRED score of 30. Duplicated reads were removed with FastUniq v1.1 (Xu et al., 2012). Reads were mapped to the bait reference using bowtie2 v2.2.9, applying options *--local* and *--very-sensitive*. Kindel V0.1.4 (Constantinides and Robertson, 2017) was utilized to create consensus sequences with a minimum coverage of 5X. Consensus sequences were split into single exon contigs which were then compared to the original bait sequences with BLAT (Kent, 2002) using a minimum identity threshold of 85%. Exons were aligned and concatenated using MAFFT v7.304 (Katoh and Standley, 2013) and catfasta2phyml.pl (Nylander, 2016). There is evidence for polyploidy in *Zanthoxylum* (Guerra, 1984; Stace et al., 1993; Kiehn and Lorence 1996) and thus greater risk of paralogs in the dataset. HybPhyloMaker is not able to filter against paralogs directly but considers the most abundant sequence to be the ortholog (Fér & Schmickl, 2018). SAMtools v1.8 and BCFtools v1.8 (Li et al. 2009) were used to filter for paralogous sequences, increasing the number of heterozygous sites from four to eight due to the presence of polyploid samples in the dataset. Loci with more than 70% missing data or more than 25% missing taxa were removed from further analyses, leaving 258 of the targeted 354 genes after filtering. Phylogenetic inference was conducted with two different approaches, on the concatenated dataset and by coalescent analysis of gene trees. For the concatenated dataset a Maximum Likelihood (ML) analysis was conducted with ExaML v3.0 (Kozlov et al., 2015) as implemented in the HybPhyloMaker pipeline. Starting parsimony trees for ExaML including 100 bootstrap replicates were created with RAxML v8.2.12 (Stamatakis, 2014) under the GTR + G substitution model. For the coalescent analysis gene trees were estimated with RAxML v8.2.12 (Stamatakis, 2014). and PartitionFinder v2.1.1 (Lanfear et al., 2016) applying the GTR + G substitution model and 100 bootstrap replicates. Single-gene tree files were combined into a single multi-NEWICK file (Junier and Zdobnow, 2010) and rooted using *Tetradium* as outgroup. ASTRAL-III v5.6.1 (Mirarab et al., 2014; Zhang et al., 2018) was employed for species-tree reconstruction. Phylogenetic trees were visualized in FigTree v1.4.4 (http://tree.bio.ed.ac.uk/software/figtree/).

We analyzed species-tree discord using quartet sampling (Pease et al., 2018), with a partitioned alignment and the ExaML phylogeny as input. For our approach, the minimum likelihood differential between the best and the second-best likelihood quartet tree was set to 2, and 300 quartet replicates were performed for every branch. For each node quartet concordance (QC), quartet differential (QD) and quartet informativeness (QI) were computed. In addition, quartet fidelity (QF) values were computed per sample. QC values measure quartet concordances, QD values assess the ratio of the two possible discordant topologies. While QI values indicate the informative capacity of the dataset to resolve a respective node, QF values provide information on the amount of concordant topologies resolved when incorporating the specific taxon/specimen.

### e. Analysis of Off-target Reads

To explore the possible utility of the by-catch, unmapped reads were extracted from the individual *.sam files resulting from HybPhyloMaker (Fér & Schmickl, 2018) using Samtools v1.9 (Li et al., 2009; Supplemental Fig. S1). Unmapped reads were then assembled de-novo using the software SPAdes v3.13.2 (Bankevich et al., 2012) and k values 21, 33, 55, and 77. Contigs were annotated using BLAST (Altschul et al., 1990) and the January 2020 release of Uniprot („UniProt“, 2019) as the reference database with an e-value cut-off of 10 e-3 and reporting only the best hit. Blast hits were filtered to retain only entries with alignment lengths > 120 bp, in order to facilitate the possible design of baits during subsequent bait refinement and to remove spurious hits. Trinotate v3.2.1 (Bryant et al., 2017) was used to create annotation reports for each sample. A custom python script (https://github.com/ClaudiaPaetzold/off-target-reads.git) was used to compare annotation reports across samples. In a first step, contigs with blast hits to non-spermatophyte gene sequences were excluded as putative contaminations. We cataloged the origin of contigs representing putative contaminations into categories bacteria, fungi, insects, rodents, humans and non-spermatophyte embryophytes. For the remaining contigs annotated as spermatophyte genes, coverage across samples was assessed and filtered to a minimum of 24 samples (50% of all samples). For these contigs, blast hits per gene per sample were assessed, and if > 8 in any sample, the gene was marked as a putatively repetitive region and excluded. We elected to not filter to one blast hit per gene per sample, in order not to exclude multiple non-overlapping fragments for the same gene. For the remaining genes, sample-specific sequences were collected into per-gene *.fasta files and aligned using MAFFT 7.304 (Katoh & Standley, 2013) with a maximum of 100 iterations and the *localpair* option. Alignments were checked visually for sequences not overlapping with the remaining alignment (these were deleted) and sorted according to copy number status. Alignments, in which each sample was represented by only one sequence, were regarded as putative single-copy genes (pSCGs) in *Zanthoxylum*, the remainder as putative low-copy genes (pLCGs). A custom python script (https://github.com/ClaudiaPaetzold/off-target-reads.git) was used to remove columns containing only gaps and trim alignment ends to a sequence coverage of 75%. Trimmed pSCG alignments were concatenated using the AMAS (Borowiec, 2016) python suite. RAxML v8.2.8 (Stamatakis, 2014) was used for ML-based tree inference on the concatenated pSCG alignment with the substitution model GTR + G. Statistical support was assessed with 1000 bootstrap replicates. In addition, gene trees of pSCG alignments were estimated with RAxML v8.2.8 and summarized in a coalescence framework using ASTRAL III v5.6.1 (Zhang et al., 2018). The resulting species trees were also subjected to quartet sampling (Pease et al., 2018) to further assess support and results were compared to those from on-target reads. Alignments of pLCGs were also visually checked and edited as necessary. Gene trees resulting from pLCGs were screened for taxon composition and amount of duplicated taxa. We differentiated between pLCG alignments in which only one or few specimens were represented by more than one sequence (pLCG_few), and putative gene families with multiple copies in all or nearly all samples (pLCG_most). As a case study their potential to provide additional information for the Hawaiian *Zanthoxylum* species was assessed. The entire workflow (Supplemental Fig. S1) and all custom python scripts are available on github https://github.com/ClaudiaPaetzold/off-target-reads.git.

## Results

### a. Raw Data and Processing of On-Target reads

The Illumina HiSeq run resulted in an average of 11,343,488 raw reads (3,851,028 - 24,070,974; Supplemental Tab. S2). After quality trimming and deduplication 62.11% of the reads remained. Of those, an average of 49.1% (29.43% to 65.89%) per sample mapped to the bait reference. Out of the 354 genes, 96 did not meet the thresholds of < 70% missing data and < 25% missing taxa, resulting in a final dataset of 258 genes. Of these 258 gene alignments, 231 included all 48 taxa and eleven included 47 taxa. The remaining 16 gene alignments contained between 36 and 46 taxa. The individual alignments ranged from 118 to 3294 bp in length and the concatenated alignment of all 258 genes sums up to 187,686 bp in length. The percentage of missing data in the individual alignments ranged from 0 to 69.1% with an average of 10.8%. The individual alignments contained between 10 and 1030 variable sites each, of which six to 498 were parsimony informative. The concatenated alignment contained 49,993 variable sites 24,030 of which were parsimony informative.

### b. Phylogenetic Analyses

The three major *Zanthoxylum* clades identified recently (Appelhans et al., 2018) are confirmed based on the analyses of the concatenated (Fig. 1) and coalescent datasets (Fig. 2), and a fourth clade has emerged comprising *Z. asiaticum* only. Clade 1 comprises all sampled *Zanthoxylum* species endemic to continental Africa, Madagascar and Mauritius (100% bootstrap support [BS, concatenated analysis, Fig. 1] /1.00 local posterior probability [lPP, coalescent analysis, Fig. 2]). It is resolved as sister to the remainder of the genus (100% BS/ 1.00 lPP) that consists of the major Clades 3 and 4, with *Z. asiaticum* (Clade 2) as sister to Clade 3 and Clade 4. The split between Clade 3 and Clade 4 is resolved with moderate support (79% BS/ 0.68 lPP). Clade 3 comprises species from continental Asia, Malesia and Australia, with a monophyletic Pacific group embedded within (100% BS/ 0.67 lPP). Clade 4 comprises species distributed across the Americas, with a second monophyletic Asian lineage and a species from the Juan Fernandéz Islands (Chile, South Pacific) embedded within (100% BS/1.00 lPP). Several backbone nodes of Clade 4 are not well supported in the coalescent species-tree (Fig. 2) but all except one node received BS values of > 90 in the concatenated analysis (Fig. 1). Support of more recent nodes is strong in either analysis. The species tree topologies from the concatenated and coalescent datasets are congruent except for three cases. Within Clade 1, *Z. ovatifoliolatum* is nested within a paraphyletic *Z. chalybeum* with low support in the concatenated analysis (Fig. 1; 58% BS). Both species are resolved as sisters in the coalescent analysis with maximum support (Fig. 2; 1.00 lPP). In the concatenated analysis, *Z. dissitum* and *Z. scandens* form a clade (Fig. 1; 75% BS) that is sister to *Z. echinocarpum*. In the coalescent analysis, *Z. dissitum* and *Z*. *echinocarpum* are resolved as sisters, with low support (Fig. 2; 0.56 lPP). *Zanthoxylum caribaeum* ssp*. caribaeum* is resolved as sister to the American clade that contains all species from sections *Pterota* and *Tobinia* within Clade 4 in the concatenated analysis (Fig. 1; 93% BS). In the coalescent analysis it is not resolved as sister to sections *Pterota* and *Tobinia*, but as an early-branching clade of Clade 4 in a part of the tree with low support (Fig. 2; 0.76 lPP). None of these cases represent a hard conflict, since at least one reconstruction method did not succeed in resolving the respective branch with high support. Concerning the deeper nodes, the phylogeny based on the concatenated dataset (Fig. 1) lacks support only at the ancestral node of Clade 3 and Clade 4 (79% BS) and at the ancestral node of American and Asian *Zanthoxylum* species within Clade 4 (85% BS). In contrast, the coalescent species tree (Fig. 2) shows a largely unsupported backbone regarding Clades 3 and 4. Both, however, show generally strongly supported younger nodes.

**Fig. 1.**
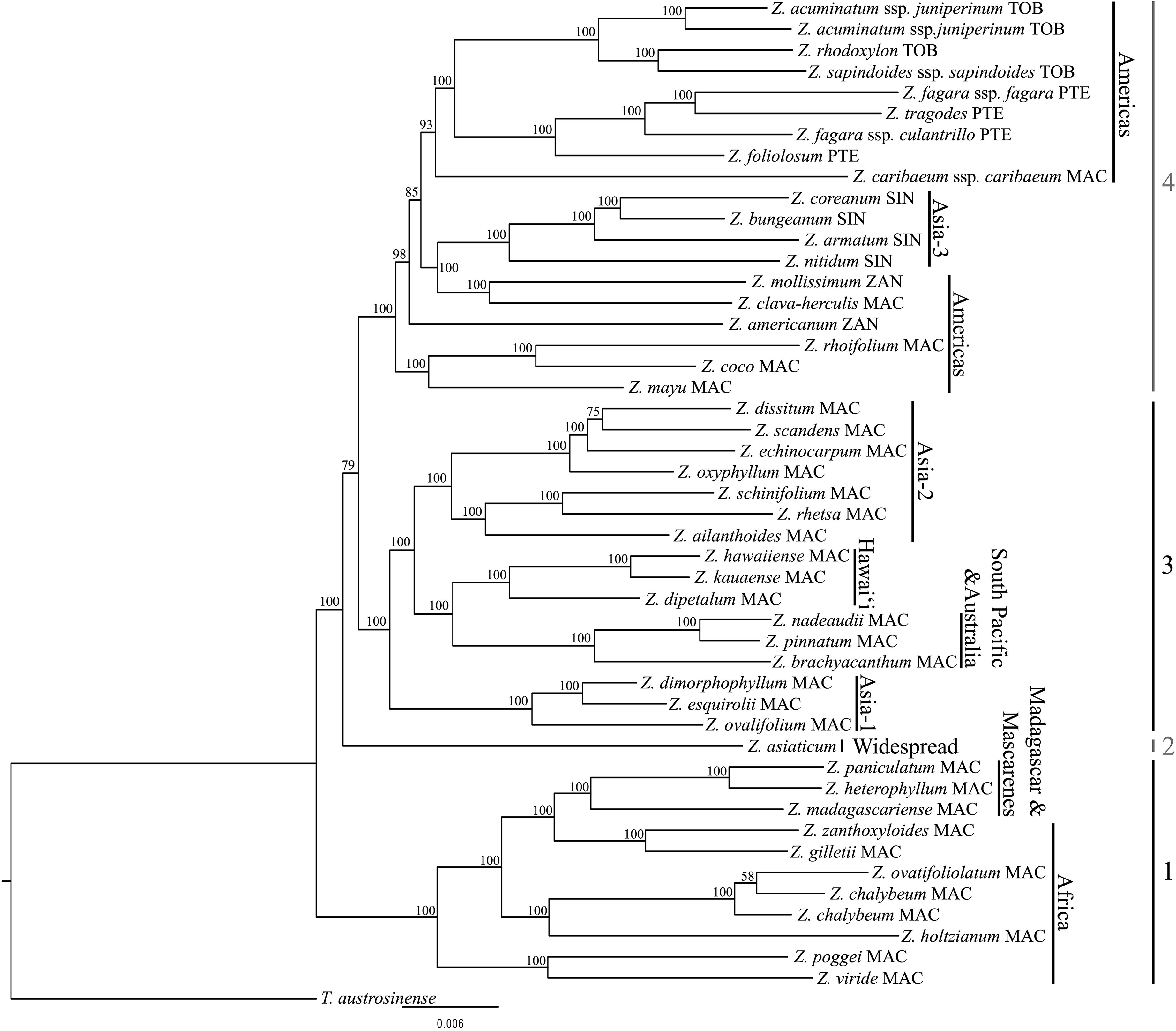
ExaML phylogenetic tree of *Zanthoxylum* based on the concatenated alignment of 258 targeted nuclear genes. Bootstrap values (BS) are shown at each branch. Abbreviations after species names refer to their current sectional classification according to Reynel (2017). MAC = *Macqueria*, PTE = *Pterota*, SIN = *Sinensis*, TOB = *Tobinia*, ZAN = *Zanthoxylum*.

**Fig. 2.**
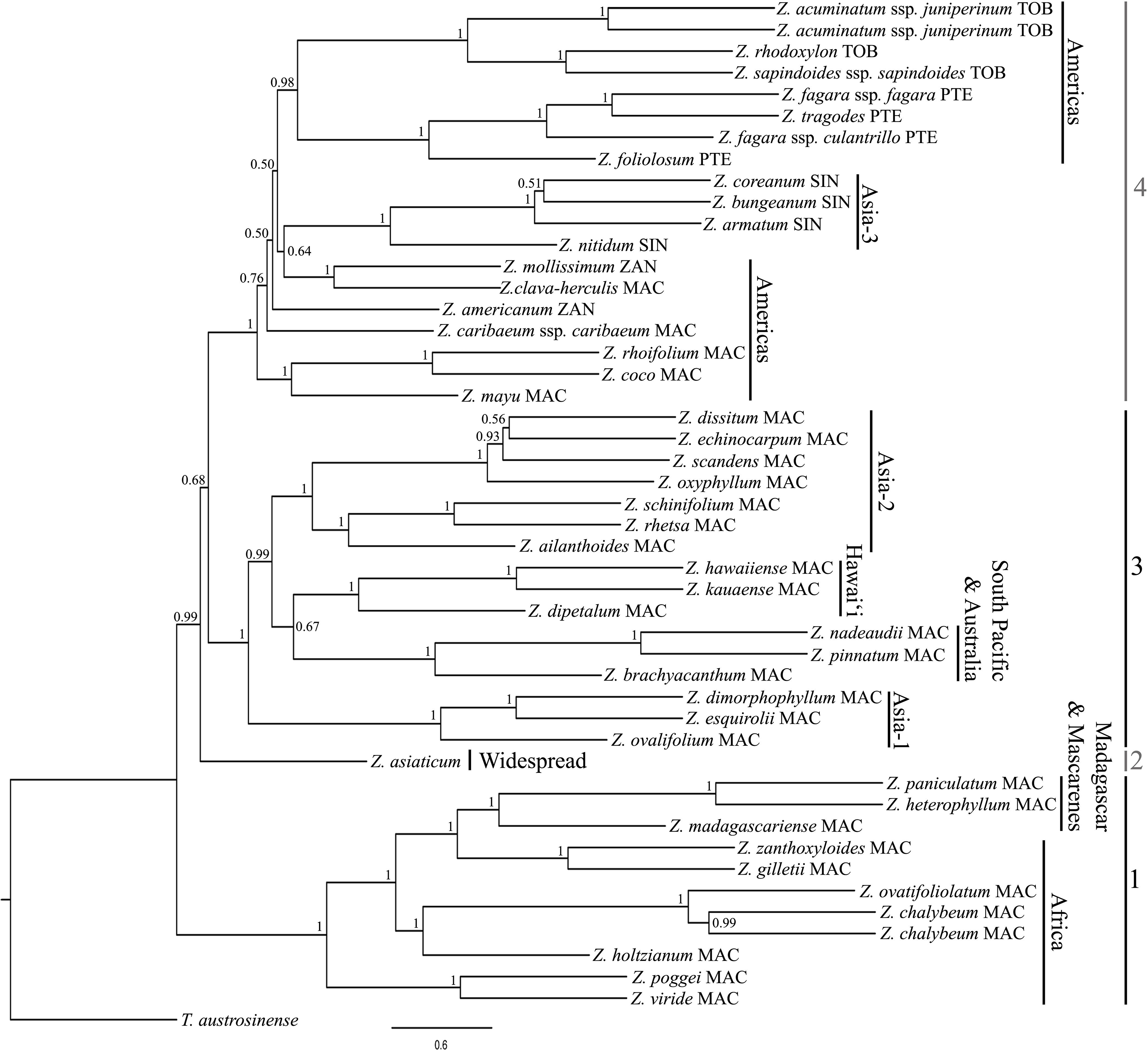
ASTRAL species tree of *Zanthoxylum* based on 258 targeted genes. ASTRAL local posterior probability (lPP) values are displayed at each branch. Abbreviations after species names refer to their current sectional classification according to Reynel (2017). MAC = *Macqueria*, PTE = *Pterota*, SIN = *Sinensis*, TOB = *Tobinia*, ZAN = *Zanthoxylum*.

### c. Sectional Relationships

Only two or three of the *Zanthoxylum* sections according to Reynel (2017) are resolved as monophyletic here. Section *Pterota* and *Tobinia* each form a monophyletic group with high support in both concatenated and coalescent analyses (Figs. 1, 2). Section *Sinensis* is only monophyletic if *Z. dimorphophyllum* is excluded. This species produces both homo- and heterochlamydeous flowers (Zhang et al., 2008). It has not been assigned to a section yet, but floral morphology places it in either section *Sinensis* (homochlamydeous) or *Macqueria* (heterochlamydeous). Section *Zanthoxylum* is polyphyletic. Of the two sampled species, *Z*. *americanum* is sister to a large part of Clade 4, while *Z*. *mollissimum* is resolved as sister to *Z*. *clava*-*herculis* of section *Macqueria*. Section *Macqueria* is largely polyphyletic and its members are scattered in all main clades. *Zanthoxylum* s.str. (= sections *Sinensis* and *Zanthoxylum*) is deeply nested within *Fagara* (= all other sections) and it is not monophyletic due to the placement of *Z. americanum*, which makes *Zanthoxylum* s.str. paraphyletic with respect to sections *Pterota* and *Tobinia* as well as *Z. caribaeum*.

### d. Concordance Analysis

A quartet concordance analysis was conducted for the main dataset with the alignment partitioned by genes and the concatenated ML tree as input (Pease et al., 2018; Fig. 3). Nodes with lower QC values are more predominant in the backbone region while the majority of the remaining nodes are strongly concordant. A consistently high QI is inferred (0.78 – 1.0) over the entire topology, as well as high QF scores for all samples (0.75 – 1.0). The split of *Z. asiaticum* as well as the divergences of Clade 3 and Clade 4 are associated with a low QD and a medium QC. The ancestor to Clades 3 and Clade 4 is characterized by a combination of low QC and high QD values. The most recent common ancestor of the Pacific lineage shows a low QC in combination with a QD value of 0. The backbone of the American-Asian lineage within Clade 4 shows several nodes with low QC values combined with medium to low QD values and short branch lengths. In two cases of topological incongruences between the concatenated and the coalescent trees, the concatenated tree showed low bootstrap support (see Results section b; placement of *Z. dissitum*, *Z. echinocarpum* and *Z. scandens*; placement of *Z. chalybeum* and *Z. ovatifoliolatum*). The respective nodes show low or medium QC values and mixed QD values.

**Fig. 3.**
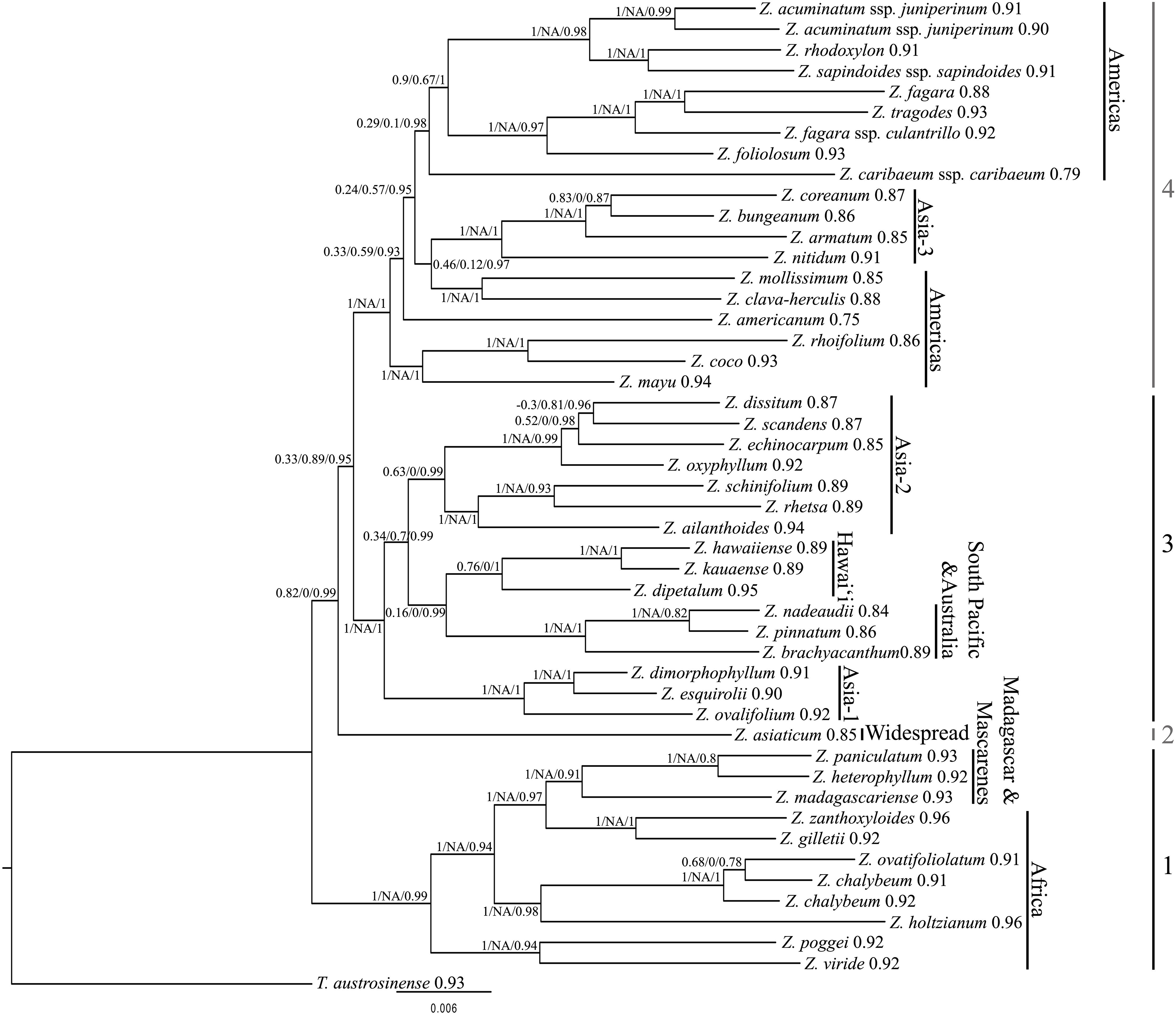
*Zanthoxylum* quartet sampling scores (300 replicates) on the basis of the ExaML tree topology. On each node Quartet Concordance (QC)/Quartet Differential (QD)/Quartet Informativeness (QI) scores are displayed. Quartet Fidelity (QF) scores are shown next to the species names.

### e. Off-Target Reads

Target enrichment sequencing resulted in an average of 3,524,882 off-target reads (1,153,183 – 8,409,705; Supplemental Tab. S2) per sample. On average, 16.95% of off-target reads (6.50 – 54.66%) were identified as putative contaminations of the plant material (Supplemental Tab. S3; Supplemental Fig. S2). The most prominent taxonomic group among the putative contaminants was fungi (531 contigs on average), followed by insects (501 contigs on average). Specimens showed major variation in their putative contamination patterns (Supplemental Tab. S3; Supplemental Fig. S2). For example, fungi represent the largest percentage of putative contaminations in the *Z. mayu* specimen, while the highest number of contigs representing putative contaminations in the *Z. chalybeum* and *Z. viride* specimens are insects and bacteria, respectively. The *Z. dimorphophyllum* and *Z. mayu* samples were taken from herbarium specimens collected in 1930 and 1955, respectively. The percentage of putative contamination in the *Z. dimorphophyllum* sample ranks among the lowest in the whole dataset, and the composition of taxonomic groups among the putative contaminants is highly similar to the average. In contrast, the *Z. mayu* sample is the only sample that shows more than 50% putative contaminants among the off-target contigs, and 74.5% of them are of fungal origin (Supplemental Tab. S3; Supplemental Fig. S2). In total, 97 pSCGs alignments with a total length of 60,521 bp could be assembled, with four of these from the chloroplast genome and another four from the mitochondrial genome (Supplemental Tab. S4). The concatenated ML tree (Fig. 4) based on these 97 pSCGs is largely congruent with the phylogenies based on targeted loci. However, node support is overall slightly lower (Figs. 1, 2, 4). Like in the coalescent analysis of the on-target reads, the coalescent pSCG tree (Supplemental Fig. S3) is characterized by lower lPP values. It shows a major deviation from both on-target phylogenetic trees (Figs. 1, 2) and the concatenated pSCGs (Fig. 4) since it does not resolve the Hawaiian *Zanthoxylum* species as sister to the South Pacific and Australian lineage. In contrast to the on-target trees, *Z. chalybeum* is monophyletic in the concatenated pSCG tree (76% BS) but polyphyletic in the coalescent pSCG tree albeit with low support (0.46 lPP). Both the concatenated and coalescent pSCG topologies diverge from on-target trees with *Z. echinocarpum* and *Z. scandens* as sister to each other (97% BS; 0.82 lPP), and *Z. dissitum* at the base to these (0.63% BS; 0.29 lPP). *Zanthoxylum mayu* is resolved as sister to *Z. coco* and *Z. rhoifolium* with high support (100% BS; 1.00 lPP) in the on-target trees but resolved as sister to the section *Pterota* and *Tobinia* clade in the coalescent pSCG tree (0.85 lPP). In the concatenated pSCGs tree its placement next to a clade with sections *Sinensis* and *Zanthoxylum* is not supported (0.24% BS). The concatenated pSCG tree diverges from all other trees as it resolves *Z. americanum* as direct sister to *Z. clava-herculis* and *Z. mollissimum* (0.83% BS; Fig. 4). Partitioned quartet sampling of the coalescent pSCG gene tree results in overall lower QC and mixed QD values (not shown) in comparison to the quartet sampling analysis of targeted data. Likewise, QI (0.11 – 0.73) and QF values (0.18 – 0.41) are generally low in the coalescent pSCG tree.

**Fig. 4.**
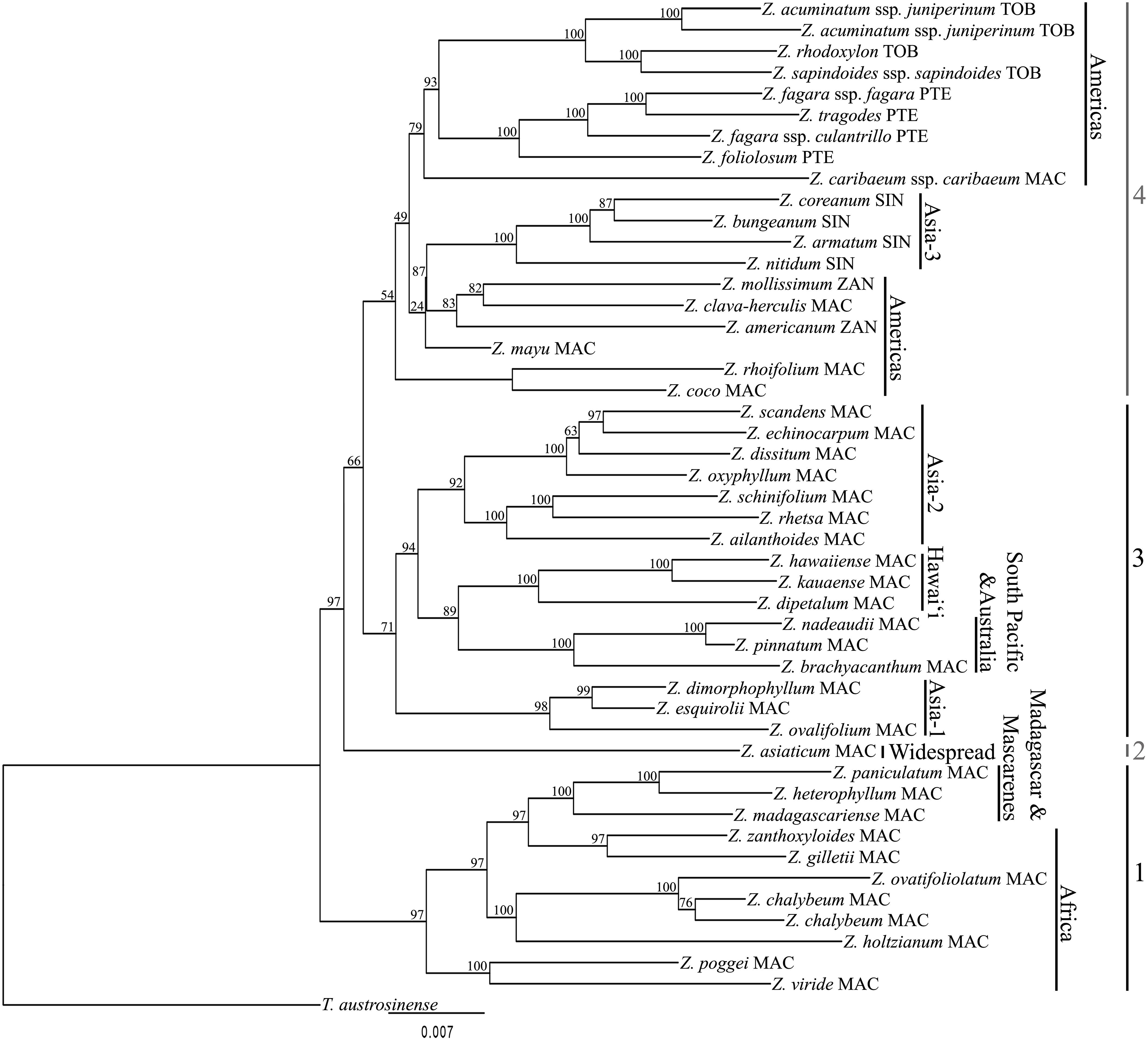
ExaML phylogenetic tree of *Zanthoxylum* based on a concatenated alignment of 97 off-target putative single-copy genes. Bootstrap-values (BS) are shown on every branch. Abbreviations after species names refer to their current sectional classification according to Reynel (2017). MAC = *Macqueria*, PTE = *Pterota*, SIN = *Sinensis*, TOB = *Tobinia*, ZAN = *Zanthoxylum*.

In total, 260 alignments based on the off-target reads were of good quality, but had at least one sample represented by more than one sequence (Supplemental Fig. S4). The gene trees inferred from these alignments were generally not well resolved. In 41 cases, only one or few specimens were represented by more than one sequence (pLCG_few), while 219 alignments constituted putative gene families with multiple copies in all samples (pLCG_most). In the 41 pLCG_few alignments, 21 to 43 (average: 31.4; standard deviation: 6.3) of the 48 samples were included and one to 15 specimens (average: 3.8; standard deviation: 3.4) were represented by two to four sequences. Some specimens were nearly always represented in the 41 pLCG_few alignments (*Z. dipetalum*, *Z. hawaiiense* [38x each], *Z. acuminatum* ssp. *juniperinum* 2, *Z. rhetsa*, *Z. rhoifolium* [37x each]), while three samples were present only in less than 10 alignments (*Z. poggei* [8x], *Tetradium austrosinense* [7x], *Z. mayu* [1x]). Nine specimens never showed any duplicated sequences and another 12 specimens were duplicated in only one or two alignments. Only two specimens had duplicated sequences in more than 10 alignments (*Z. asiaticum*, *Z. rhoifolium* [12x each]). As a case study for the informative value of these gene trees, we focused on the Hawaiian lineage. In 11 of the 41 pLCG_few alignments, one or more Hawaiian species were represented by more than one sequence and in four of these alignments, only Hawaiian species had duplicated sequences. In 18 out of the 41 alignments, the gene trees resolved the Hawaiian lineage as polyphyletic with at least low support. In some cases, the relationships of the polyphyletic Hawaiian groups could not be determined due to low resolution of the gene trees. In case of reasonable resolution of gene trees, in all cases of a polyphyletic Hawaiian group, one copy resolved Hawaiian species as closely related to South Pacific and Australian species (*Z. brachyacanthum*, *Z. pinnatum*, *Z. nadeaudii*), while the other copy was most closely related to Asian species, most frequently to the clade of *Z. ailanthoides*, *Z. rhetsa* and *Z. schinifolium* (Supplemental Fig. S4). This latter relationship of the Hawaiian lineage was also found in the coalescent analysis of the pSCGs (Supplemental Fig. S3).

## Discussion

### a. Phylogenetic Relationships

We present the first phylogenomic study for the genus *Zanthoxylum* using a target enrichment high throughput sequencing approach. Concatenated and coalescent analyses of both the on-target as well as the off-target pSCG dataset resulted in phylogenetic trees largely congruent with the recently published *Zanthoxylum* phylogeny based on four genes only (Appelhans et al., 2018). However, phylogenetic resolution and support in several clades have greatly improved in this study. The observation that several nodes especially in the backbone of the phylogeny still received mixed or low support provides evidence that low support is not only due to the limited size of the Sanger dataset but to conflicts in the dataset that may be attributed to reticulate evolution (Figs. 1, 2; Appelhans et al., 2018). Herein, the genus *Zanthoxylum* is divided into four major clades.

#### The African clade (Clade 1)

*Zanthoxylum* species endemic to the African continent, Madagascar and the Mascarene Islands represent a monophyletic lineage that is sister to the remainder of the genus. Within this clade, the accessions from Madagascar and the Mascarene Islands are resolved as monophyletic (Clade 1; Fig. 1, 2). The West and Central African *Z. lemairei* was nested within the Malagasy clade in the study by Appelhans et al. (2018), but with low support. This species was not sampled in the present study, so it remains unclear whether the Malagasy and Mascarene species represent a monophyletic lineage that evolved from a single colonization event from mainland Africa, or if there were either two colonization events or a single colonization event with one back-dispersal to the African mainland. *Zanthoxylum heterophyllum* and *Z. paniculatum* are the only *Zanthoxylum* species from the Mascarenes and *Z. paniculatum* is an extremely rare endemic to the island of Rodrigues (Bone, 2004). The two species are resolved as sisters in our study and they might have evolved from a common Malagasy ancestor. A larger taxon sampling of Malagasy species is needed to draw further conclusions about the relationships of these two species with regards to the Malagasy species. With the exception of *Z. chalybeum* and *Z. ovatifoliolatum*, the relationships in Clade 1 are well resolved and quartet sampling revealed no signals of reticulate evolution in this clade. The on-target concatenated analysis (Fig. 1) and off-target coalescent pSCG tree (Supplemental Fig. S3) both resolved *Z. chalybeum* as paraphyletic with respect to *Z. ovatifoliolatum*, but with low bootstrap support (58% BS; 0.46 lPP) and the quartet sampling showed a low QC value and a QD value of 0 (Fig. 3). However, the on-target coalescent tree (Fig. 2) and concatenated pSCG tree (Fig. 4) resolved the two specimens of *Z. chalybeum* as monophyletic and sister to *Z. ovatifoliolatum* with medium to high support, so that this topology is considered more likely.

#### The *Zanthoxylum asiaticum* clade (Clade 2)

*Zanthoxylum asiaticum*, formerly recognized as *Toddalia asiatica*, represents a separate lineage and is sister to the major Clades 3 and 4 in the analyses of the on-target dataset with high support. The off-target pSCGs resolved the species as sister to Clade 4, although the relationship had moderate support and low QC and QD values in quartet sampling. In a previous study the identical *Z. asiaticum* individual was resolved as sister to the African species (Clade 1 in the present study; Appelhans et al., 2018). Appelhans et al. (2018) sampled several outgroup taxa including most species of *Fagaropsis*, *Phellodendron* and *Tetradium*, as well as the more distantly related *Acronychia* and *Melicope*.

#### The Asian – Pacific – Australian clade (Clade 3)

Clade 3 consists of four subclades: the Asia-1 subclade, the South Pacific-Australian subclade, the Hawaiian subclade, and the Asia-2 subclade. Resolution, support and quartet sampling values are high in Clade 3 except for two nodes regarding the Pacific lineages and the relationships among *Z. dissitum*, *Z. echinocarpum* and *Z. scandens* (Figs. 1-4). Three out of four Hawaiian species are sampled here, and the herein missing *Z. oahuense* was sampled by Appelhans et al. (2014). It showed a close relationship with Z*. hawaiiense* and *Z. kauaense*, so the Hawaiian species are most likely monophyletic. *Zanthoxylum dipetalum* was confirmed as the earliest diverging lineage and sister to the remaining Hawaiian species (Fig.1, Fig., 3, Appelhans et al., 2014), which is supported by distinct morphological features (Hillebrand, 1888; Wagner et al., 1999) such as (usually) two petals, lowest pair of leaflets reduced in size, and larger fruits with a beaked apex. The sister group of the Hawaiian clade comprises the Australian *Z. brachyacanthum*, and two Pacific species, *Z. nadeaudii* endemic to the Society Islands and *Z. pinnatum,* which is more widespread in the South Pacific (Lord Howe Island to the Austral Islands; Butaud and Meyer, 2004). These two small lineages are sister to an Asian lineage (Asia-2 subclade) with species ranging from Southeast Asia to China (Figs. 1, 2; Clade 3). In the most recent *Zanthoxylum* phylogeny (Appelhans et al., 2018) this Pacific group was resolved as monophyletic based on the plastid genes *trn*L-*trn*F and *rps*16 data but polyphyletic based on nuclear ITS and ETS sequences. Thus, the authors hypothesized a hybridization event prior to the colonization of the islands. Herein, we analyzed gene tree discordance using quartet sampling (Pease et al. 2018). For the ancestral Pacific node the QC value (0.16; Fig. 3) indicates a strong discord with one dominant discordant topology (QD = 0), supporting the hypothesis of a previous hybridization event. Furthermore, it is striking that 18 out of the 41 off-target pLCG_few alignments resolved the Hawaiian lineage as polyphyletic (Supplemental Fig. S4). In nearly all of these cases, one Hawaiian gene lineage was resolved as sister to South Pacific and Australian species, while the other gene lineage was closely related to an Asian clade (Asia-2) and the close relationship to the Asia-2 clade was also found in the coalescent pSCG tree (Supplemental Fig. S3). Chromosome numbers are known only for one of the four Hawaiian species, *Zanthoxylum hawaiiense*, where a chromosome count of 136-144 suggests an octoploid cytotype (Kiehn & Lorence, 1995). The majority of *Zanthoxylum* species appear to be tetraploids with 68 to 72 chromosomes (e.g. Guerra, 1984; Zhang et al., 2008), so that the Hawaiian lineage might be the result of an allopolyploidization event prior to the colonization of the Hawaiian Islands. Further cytological data of Hawaiian, South Pacific and Asian taxa will be crucial to provide insights into the ploidy levels and evolutionary relationships within these lineages.

In subclade Asia-1 (comprising *Z. dimorphophyllum*, *Z. esquirolii* and *Z. ovalifolium*), both concatenated and coalescent analyses show high support for the clade (100% BS, 1.00 lPP; Figs. 1-2) and there are no conflicting quartet topologies (Fig. 3). *Zanthoxylum esquirolii* and *Z. dimorphophyllum*, both common in South China, are resolved as sister to each other. However, *Zanthoxylum dimorphophyllum* is also native to other regions from Central China to Thailand and Vietnam (Zhang et al., 2008). *Zanthoxylum ovalifolium* is absent from China (Zhang et al., 2008; was mistakenly synonymized with *Z. dimorphophyllum* previously) but shows a broad distributional range from the Himalayas and India to Australia (Hartley 2013).

Within the Asia-2 subclade, the relationships among *Z. dissitum*, *Z. echinocarpum* and *Z. scandens* from southern China are resolved incongruently in our analyses with high to moderate support (Figs. 1 – 4). These three species formed a clade in Appelhans et al. (2018) as well, with an identical topology as in the concatenated analysis of the on-target reads in the present study. QC values are low or even negative for the respective nodes (Figs. 3, 4) and QD values are relatively high (0.81 to 0.89 in Figs. 3, 4), suggesting incomplete lineage sorting as the likely cause for the discord. Several species including *Z. laetum* and *Z. calcicola* have been hypothesized to be close relatives of the morphologically variable *Z. scandens* (Hartley, 1966). These species, the several morphotypes of *Z. scandens* as well as the varieties of *Z. dissitum* and *Z. echinocarpum* (Zhang et al., 2008) need to be included in future analysis in order to better understand relationships in this lineage.

#### The American-eastern Asian clade (Clade 4)

Clade 4 includes all American species and a subclade of species from eastern Asia (Asia-3 subclade or section *Sinensis*). This biogeographically disjunct clade (also see Valcárcel and Wen, 2019) is morphologically diverse and includes taxa that represent all five sections recognized by Reynel (2017). The concatenated analysis of the on-target reads resolved the backbone phylogeny of Clade 4 well (one node with <90% BS; Fig. 1), but branch lengths are generally short. The coalescent tree and the off-target pSCGs results show lower support similar to the results of Appelhans et al. (2018; Figs. 2, 4). Quartet sampling reveals strong quartet discordance (low QC values) in combination with high QD values, with one exception (QD = 0.1) (Fig. 3). Combined with the consistently short branches in the backbone phylogeny of Clade 4 this pattern gives an indication for incomplete lineage sorting during rapid radiation in the past.

Our analyses reveal that *Z. clava-herculis*, distributed in the southern USA and northern Mexico, and the Central American *Z. mollissimum* are the closest relatives of the Asian lineage of Clade 4, although strong support for this placement is only apparent in the concatenated analysis of the on-target reads. A close relationship to more temperate or subtropical American *Zanthoxylum* species is also indicated by the ecology of the embedded Asian lineage (Asia-3 subclade; section *Sinensis*) as most of their members are well adapted to a temperate and subtropical climate in Asia, and only *Z. nitidum* is also present in tropical Asia (Zhang et al., 2008).

Only 15 of the several dozen members of South and Central American and Caribbean species are sampled, so detailed hypotheses about their relationships will not be made here. Two subspecies of *Z. fagara* (ssp. *fagara* and ssp. *culantrillo*) have been sampled and they are resolved as paraphyletic with *Z. tragodes*. *Zanthoxylum fagara* is a widely distributed and morphologically diverse species. The subspecies *culantrillo* differs remarkably from the typical form by its spinulose fruit. Reynel (2017) described that intermediates between the two subspecies exist in regions where they co-occur. Our study suggests that *Zanthoxylum fagara* ssp. *culantrillo* should be regarded as a separate species from *Z. fagara* and the intermediate specimens might represent interspecific hybrids of the two species.

*Zanthoxylum mayu* is one of two *Zanthoxylum* species that occur on the Juan Fernández Islands, Chile. It is endemic to Robinson Crusoe Island (Masatierra), while the second species, *Z. externum*, is endemic to Alejandro Selkirk Island (Masafuera; Penneckamp, 2019).

Spatial/geographic isolation (between islands) has been identified as the primary driver of speciation on the Juan Fernández Islands (Stuessy et al., 1998; Stuessy, 2020), which results in species pairs with one species endemic to Robinson Crusoe Island and one species endemic to Alejandro Selkirk Island as it is the case in *Zanthoxylum*. Engler (1896) placed *Z. mayu* in the monotypic section *Mayu* (*Z. externum* had not been described then), which is morphologically only differentiated from its closest Central and South American relatives (section *Macqueria* series Paniculatae sensu Engler, 1896) by its inflorescence type, which is an axillary raceme in section *Mayu* versus panicles in section *Macqueria* series Paniculatae (Engler, 1896). Reynel (2017) did not recognize section *Mayu* and included it in section *Macqueria*, which is supported by our results.

### b. Sectional classifications

Our results reveal that several of the morphological sections recently described or newly circumscribed by Reynel (2017) are polyphyletic. The mainly Caribbean section *Tobinia*, characterized by 3-merous flowers (rarely 4-merous; Reynel, 2017), and the Neotropical section *Pterota*, which is distinguished by its winged leaf rhachis and 4-merous flowers (Reynel 2017), are both monophyletic. The temperate Asian section *Sinensis*, which has homochlamydeous flowers, might be monophyletic. Section *Zanthoxylum* is resolved as polyphyletic due to the placement of *Z*. *americanum* and *Z*. *clava*-*herculis*. The position of *Z. americanum* could not be clarified in this study. Appelhans et al. (2018) resolved it as sister to the remainder of section *Zanthoxylum* and section *Sinensis*. If a wider taxon sampling in future studies confirms this, section *Zanthoxylum* would form a grade with section *Sinensis* nested within it. *Zanthoxylum clava-herculis* was placed in section *Zanthoxylum* (Engler, 1896), and Reynel (2017) moved the species to section *Macqueria*, but highlighted that this was provisional. Like *Z*. *dimorphophyllum*, *Z. clava-herculis* exhibits characters that are transitional between the two sections. Heterochlamydeous flowers are typical for section *Macqueria*, whereas a deciduous perianth in fruiting carpels, a conspicuous dorsal gland in the ovules as well as a globose stigma are characteristic for section *Zanthoxylum* (Reynel, 2017). Section *Zanthoxylum* sensu Reynel (2017) consists of only three species and the only species missing in our study is *Z. ciliatum*. This species shares clear morphological similarities with *Z. mollissimum* (tepals apically pubescent, base of staminal filament pubescent), and their close relationship is very likely. The Asian section *Sinensis* is monophyletic only if *Z. dimorphophyllum*, a species comprising both homo- and heterochlamydeous flowers, is excluded from it. The species was originally described as a *Zanthoxylum* species, at a time, when *Fagara* was still recognized as a separate genus (Hemsley, 1895; Hartley, 1966). Accordingly, it could be considered to be part of Reynel’s (2017) section *Sinensis*. However, Engler (1896) placed the species in *Fagara* section *Macqueria*. Our study places the species in a clade with species from section *Macqueria*. Section *Macqueria* is highly polyphyletic and its members are found in all main clades, and its polyphyly has already been documented by Appelhans et al. (2018). Section *Macqueria* needs to be split up into at least four sections in order to establish a classification of monophyletic sections (Fig. 1). The type species of section *Macqueria* is *Z. heterophyllum* from the Mascarene Islands (Reynel, 2017). This species is part of Clade 1 in our analyses and the name *Macqueria* should therefore be applied to the African, Malagasy and Mascarene species of *Zanthoxylum* only. A formal proposal of a new sectional or subgeneric classification is premature at this stage, and the taxon sampling especially regarding Central and South American and Chinese species needs to be increased significantly in future studies. Nevertheless, the four major clades recognized in our study set the foundation for a subgeneric classification of *Zanthoxylum*.

### c. Off-target reads as a source of additional information

In previous studies, off-target read information has mainly been utilized for the assembly of partial or complete plastid or mitochondrial genomes (e.g., Weitemier et al., 2014; Ma et al., 2021). Here, we identified 89 off-target nuclear pSCGs in addition to four mitochondrial and four plastid pSCGs (Supplemental Tab. S4), and 260 pLCGs are recovered in at least half of all samples. The *Zanthoxylum* phylogeny inferred from the concatenated pSCGs phylogenetic tree (Fig. 4) is largely congruent with the topologies from on-target phylogenies (Figs. 1, 2). Thus, the newly identified pSCGs present a useful resource to complement the targeted dataset and may also be employed to improve the bait set for future studies. pLCGs, on the other hand, can be utilized to evaluate if a locus represents a gene family or if it only duplicated in one or several taxa. The reasons why a locus is duplicated in few taxa are manifold and duplicated loci might represent the parental lineages of a hybrid taxon. In this study, such information from pLCGs is shown to support the hypothesis of a past hybridization event prior to the colonization of the Hawaiian Islands based on the quartet concordances of the targeted data. The reasons why a specimen is missing in the alignments of a pLCG are also manifold, and a likely explanation is the low coverage of sequence reads. *Zanthoxylum mayu* is only included in a single pLCG_few alignment. It is the sample from an old herbarium specimen with the second lowest number of reads that mapped to the baits and the sample with the highest percentage of putative contamination (Supplemental Tabs. S2, S3). Hence, information from off-target pSCG and pLCG loci should be corroborated by subsequent targeted sequencing to conclusively capture sequences across samples and copy numbers. Putative contaminations identified for the off-target reads (Supplemental Tab. S3, Supplemental Fig. S2) may give an indication for the minimum sequencing coverage necessary for sufficient data when working with leaf material of which sampling or subsequent conservation treatments are unknown.

## Supplemental Data

Tab. S1. Accessions of *Zanthoxylum* and closely related genera used for the bait design.

Tab. S2. Illumina Sequencing output information.

Tab. S3. Contigs of assembled off-target reads considered as putative contamination due to best BLAST hits to non-spermatophyte genes and respective phylum of Blast hit.

Tab. S4. Functional Annotation of 97 off-target pSCGs according to UniProt. Abbreviations: At = *Arabidopsis thaliana*, Glyma = *Glycine max*, MTR = *Medicago truncatula*, Os = *Oryza sativa*.

Fig. S1. Assembly starting with read mapping to target sequences. Intermediate results including file formats in white boxes, final alignments in grey boxes. Intermediate steps in the pipeline including software or scripts used are depicted on connectors. Solid lines: off-target pipeline. Dashed lines: on-target pipeline. Grey arrows: subsequent phylogenetic analyses.

Fig. S2. Pie charts displaying the distribution of contigs assembled from putative contaminants in the off-target reads for the whole dataset and representative samples. For all shown examples, the ratio of Spermatophyte vs. non-Spermatophyte reads (= putative contamination) is shown on the left and the taxonomic composition of the non-Spermatophyte reads is shown on the right; A) average of the whole dataset; B) *Z. dipetalum* (silica material); C) *Z. chalybeum* specimen 1 (silica material); D) *Z. viride* (silica material); E) *Z. dimorphophyllum* (herbarium sample collected in 1930); F) *Z. mayu* (herbarium sample collected in 1955).

Fig. S3. ASTRAL species tree of *Zanthoxylum* based on 97 pSCGs. ASTRAL local posterior probability (lPP) values are displayed at each branch. Abbreviations after species names refer to their current sectional classification according to Reynel (2017). MAC = *Macqueria*, PTE = *Pterota*, SIN = *Sinensis*, TOB = *Tobinia*, ZAN = *Zanthoxylum*.

Fig. S4. Four representative pLCG_few trees that show gene families and an example of a polyphyletic Hawaiian lineage. Single RAxML gene trees are displayed with bootstrap values shown at branches relevant to Hawaiian species (*Z. dipetalum*, *Z. hawaiiense* and *Z. kauaense*) and their closest relatives. Colors indicate Clades 1 to 4 from Figures 1-4. A) the CB12 gene is recovered with three orthologs; B) the VQ9 gene is recovered with two orthologs; C) several specimens have duplicated sequences in the DMP5 gene. The Hawaiian lineage is resolved as monophyletic and is sister to Pacific and Australian species like in Figs. 1 and 2; D) several specimens have duplicated sequences in the gene GT643. The Hawaiian group is resolved as polyphyletic with sequences of all species (*Z. dipetalum, Z. hawaiiense* and *Z. kauaense*) resolved as sister to Pacific species and a second sequence of *Z. dipetalum* is resolved as the closest relative of the Southeast Asian *Z. rhetsa* and *Z. schinifolium*. A) and B) are midpoint-rooted, while C) and D) have been rooted at Clade 1 because the outgroup was not sampled in either case.

## Data Availability Statement

The Bait set used herein is available at (…). Raw reads for all specimens are available at the NCBI Sequence Read Archive under BioProject Number (…). Alignments and tree files are available on Dryad (…). Python scripts used for analyzing off-target reads are available on github https://github.com/ClaudiaPaetzold/off-target-reads.git.

