## Supplementary figures and images for "Target enrichment improves phylogenetic resolution in the genus *Zanthoxylum* (Rutaceae) and indicates both incomplete lineage sorting and hybridization events"

### Supplemental Figure S1

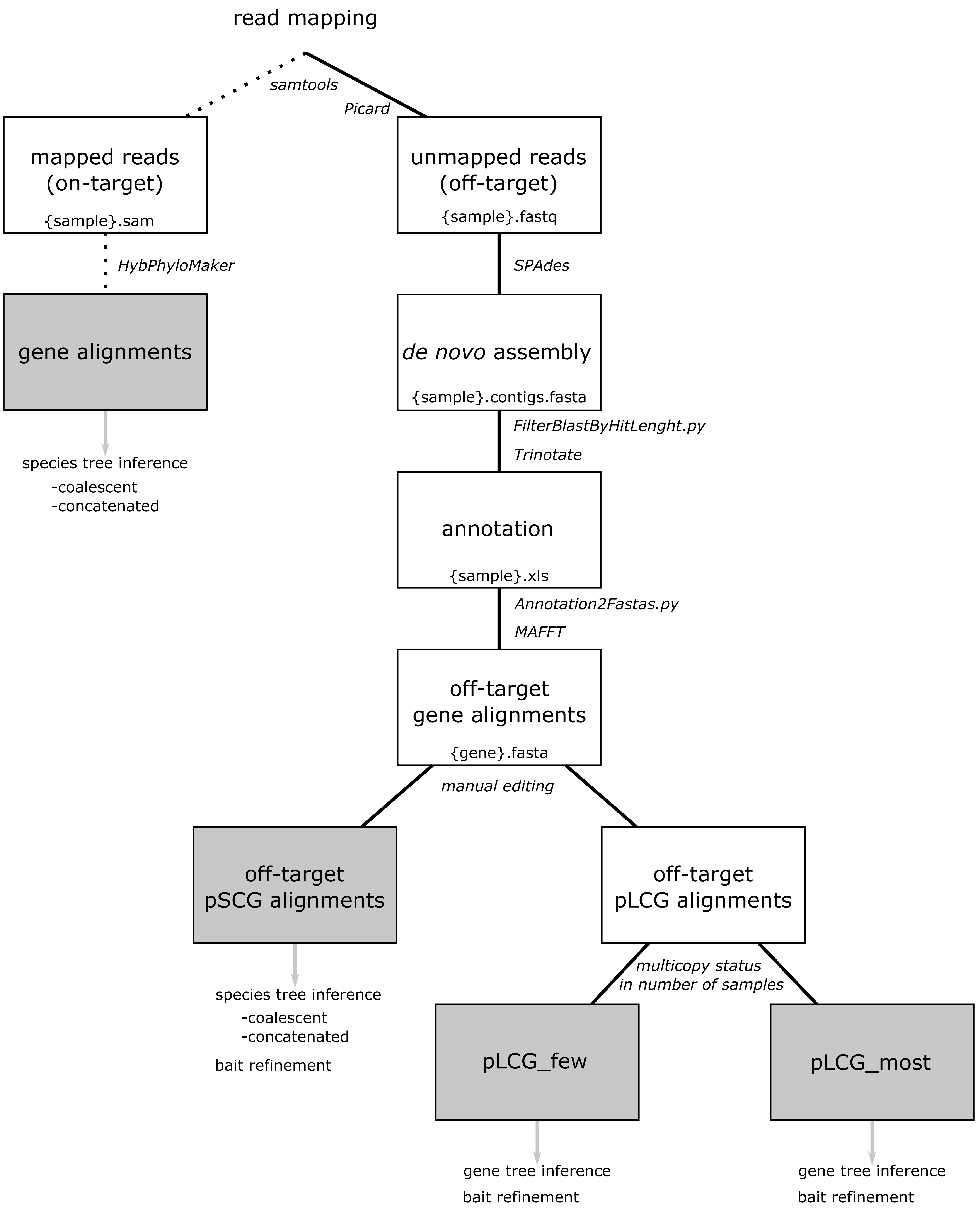

### Supplemental Figure S2

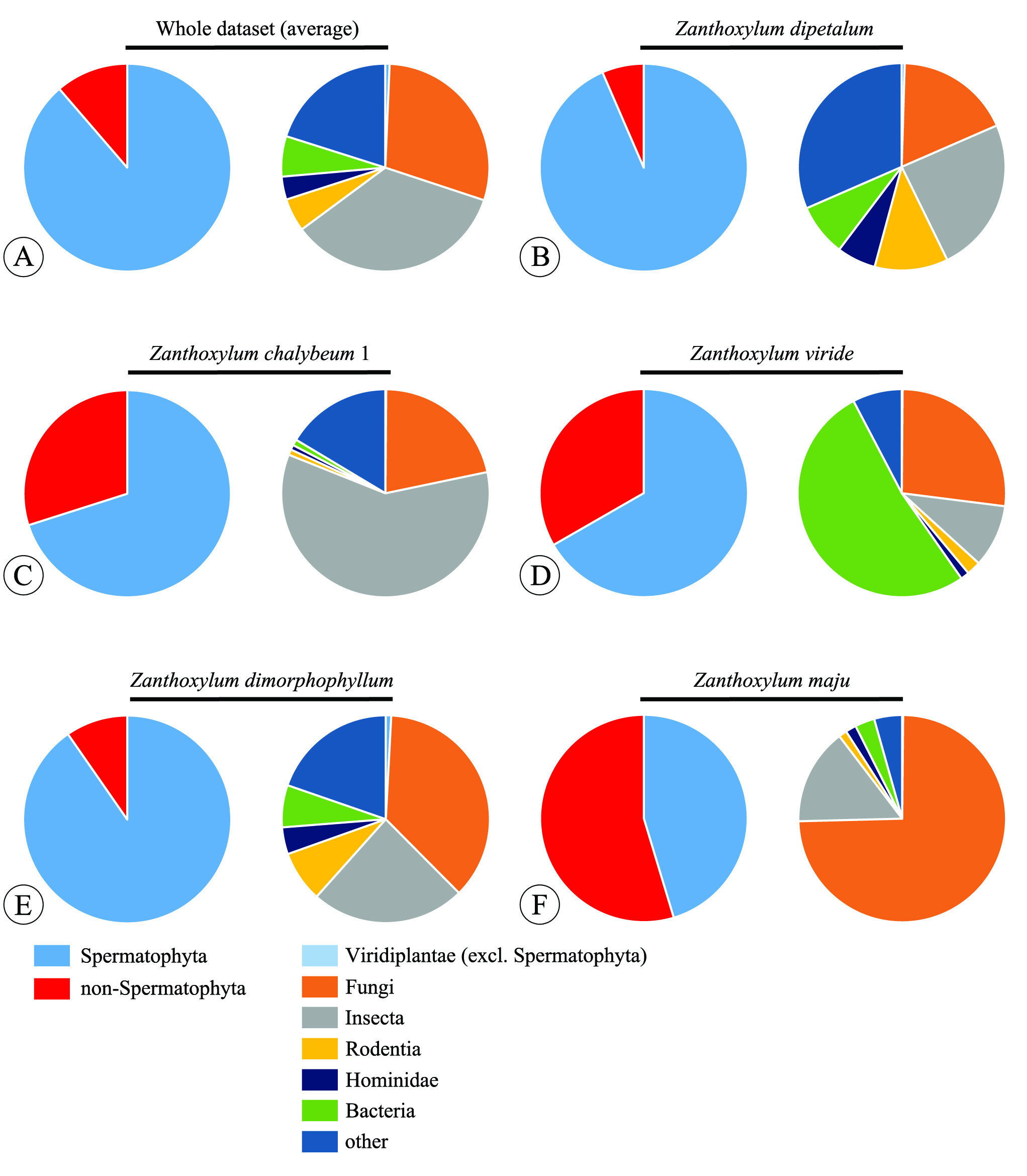

### Supplemental Figure S3

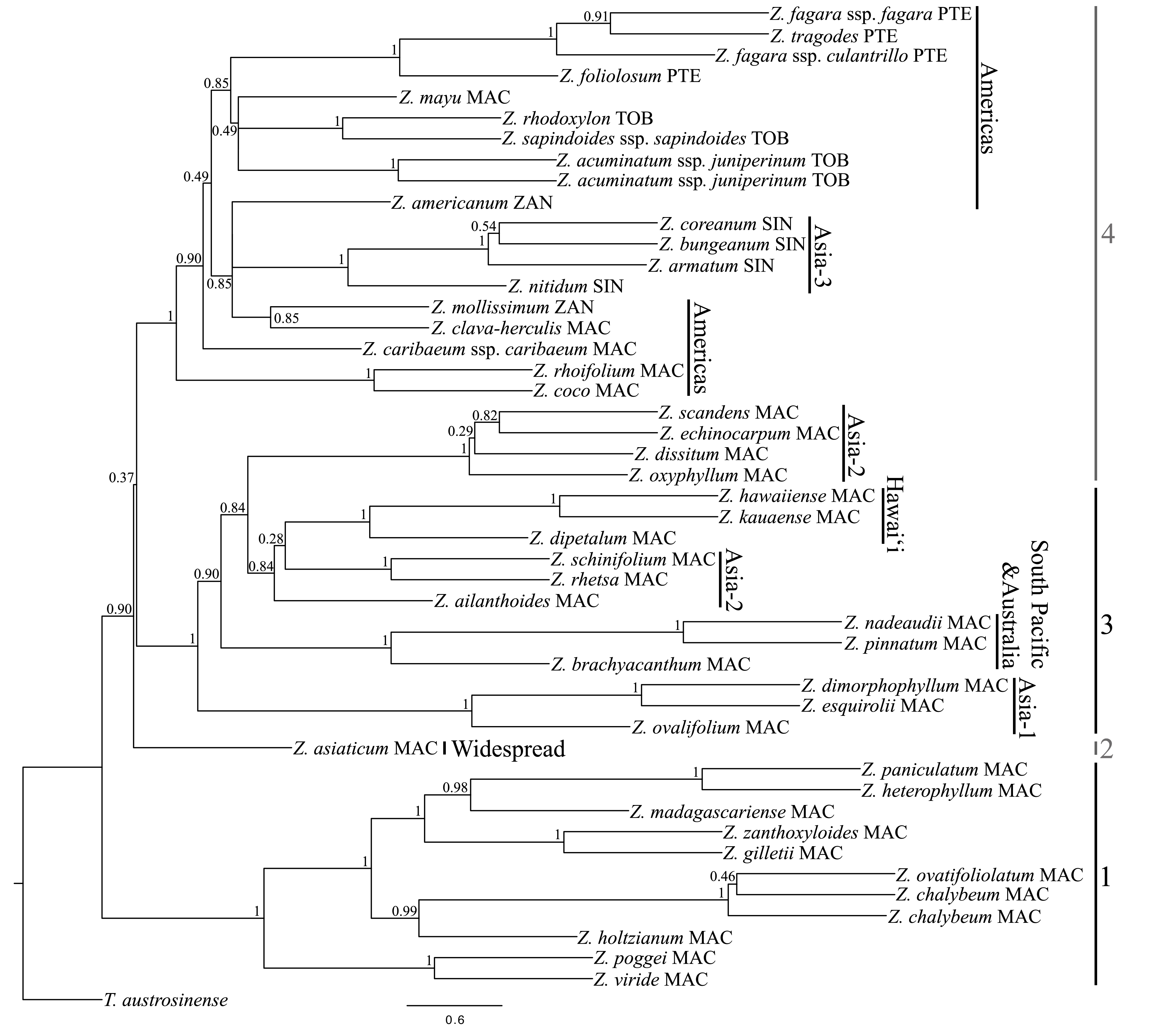

### Supplemental Figure S4

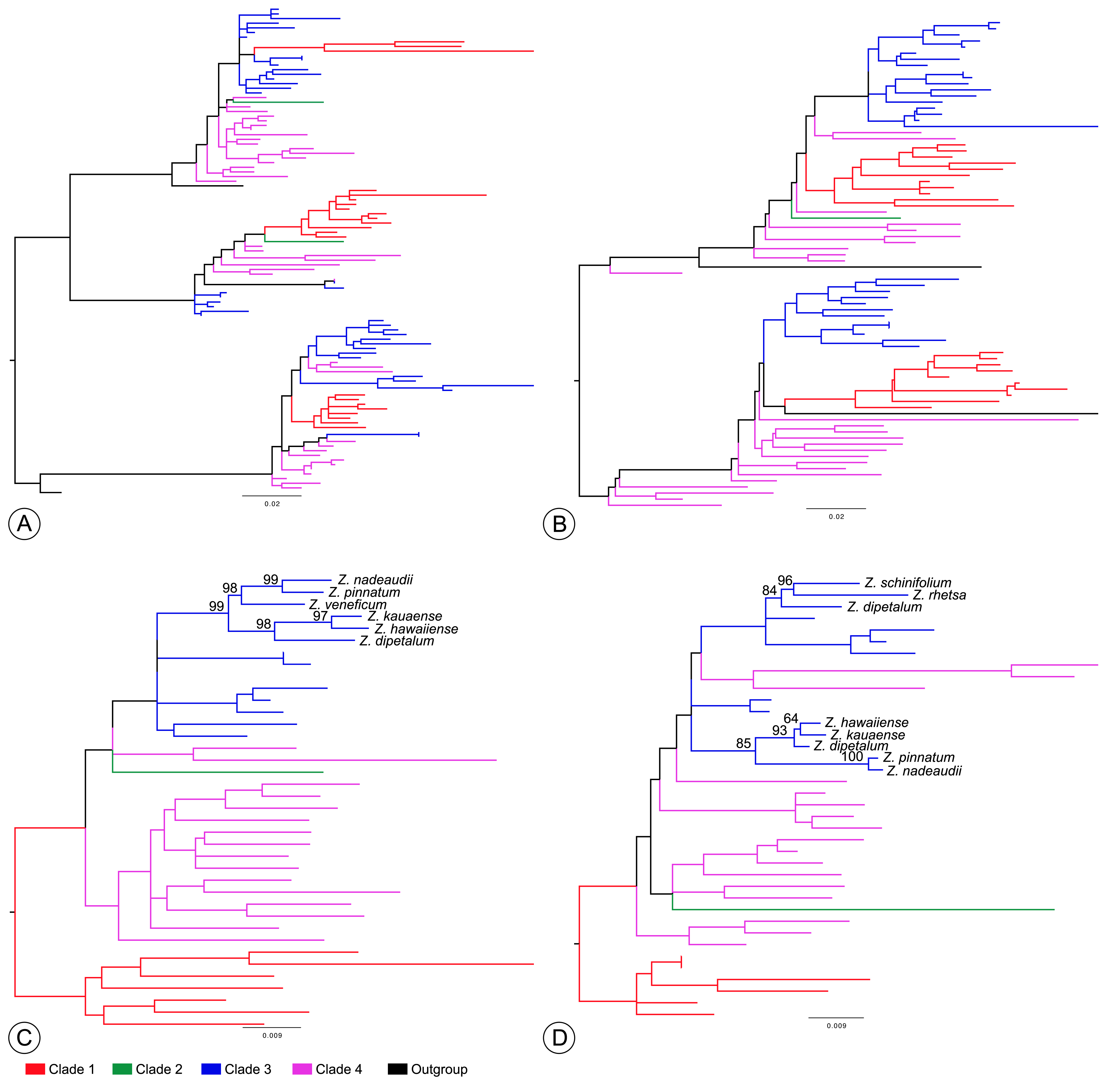
